# EEG in Motion: Using an Oddball Task to Explore Motor Interference in Active Skateboarding

**DOI:** 10.1101/2020.06.08.136960

**Authors:** Daniel Robles, Jonathan W. P. Kuziek, Nicole A. Wlasitz, Nathan T. Bartlett, Pete L. Hurd, Kyle E. Mathewson

## Abstract

Recent advancements in portable computer devices have opened new avenues in the study of human cognition outside research laboratories. This flexibility in methodology has led to the publication of several Electroencephalography (EEG) studies recording brain responses in real-world scenarios such as cycling and walking outside. In the present study, we tested the classic auditory oddball task while participants moved around an indoor running track using an electric skateboard. This novel approach allows for the study of attention in motion while virtually removing body movement. Using the skateboard auditory oddball paradigm, we found reliable and expected standard-target differences in the P3 and MMN/N2b event-related potentials (ERPs). We also recorded baseline EEG activity and found that, compared to this baseline, alpha power is attenuated in frontal and parietal regions during skateboarding. In order to explore the influence of motor interference in cognitive resources during skateboarding, we compared participants’ preferred riding stance (baseline level of riding difficulty) vs their non-preferred stance (increased level of riding difficulty). We found that an increase in riding difficulty did not modulate the P3 and tonic alpha amplitude during skateboard motion. These results suggest that increases in motor demands might not lead to reductions in cognitive resources as shown in previous literature.

## Introduction

For decades, the study of cognitive electrophysiology using Electroencephalography (EEG) has taken place inside highly controlled research facilities as EEG signals are easily contaminated by a myriad of environmental factors (Luck, 2014). EEG research has informed our understanding of human attention, yet, this knowledge generally comes from paradigms that isolate participants in faraday cages to avoid electromagnetic fields and other sources of noise that can compromise data quality (Puce & Hämäläinen, 2017). Over recent years, developments in minicomputers such as the Raspberry pi and mobile phones have allowed such studies to move outside the lab and into the real world, resulting in a growth of mobile EEG studies within ecologically rich environments (Askamp & van Putten, 2014; Kontson et al., 2015; Cruz-Garza et al., 2017; Kuziek et al., 2018).

Mobile EEG research allows us to further understand attention and brain function by exploring cognitive functions using classic paradigms in real-world scenarios. For example, an oddball task in which participants respond to rare stimuli and ignore frequent distractors has classically been used to study attentional allocation. The ERP (event-related potential) known as the P3, a positive voltage increase associated with stimulus novelty in the oddball task (Polich et al., 2003), has been widely reported in EEG studies for decades since its amplitude is modulated by increases in task effort (Luck, 2014). This robust ERP is ideal to explore brain responses in more complex real-world environments due to its large signal-to-noise ratio and replicability in the oddball task (Zamrini et al., 1991). Furthermore, the oddball paradigm allows for a dual-task approach where a primary task is presented concurrently with the oddball stimuli. These paradigms assume that resources needed to perform a primary task increase with task difficulty, and compete with cognitive resources simultaneously devoted to the oddball task, leading to measurable reductions in P3 amplitude (Kok, 2001). There is a long history of using the P3 as a measure of resource allocation in realistic tasks such as studies of pilots (Sirevaag et al., 1993), as well as video game training studies showing that P3 magnitude to a secondary task increases as a function of training in a difficult primary task (Maclin et al., 2011, Mathewson et al., 2012).

In paradigms involving motion (walking etc.), resource allocation has been investigated via cognitive-motor interference (CMI). CMI refers to the costs in cognitive and/or motor performance during dual-tasking due to the allocation of limited brain resources (Leone et al., 2017). CMI paradigms allow measuring the markers of cognitive resource allocation or motor performance during increased dual-task demands. One important assumption in dual-task paradigms is the potential involvement of common neural processes in cognitive and motor function (De Sanctis et al., 2014). PET studies have shown that activation of the lateral and superior parietal cortices, as well as prefrontal regions, is associated with the executive processes of updating, shifting, and inhibition that facilitates dual-tasking (Collette et al., 2005). Changes in the motor domain have been previously associated with increased dual-task demands during walking. Gait speed, a general index of functional performance, has been shown to decrease during dual-task interference and such a decrease is considered an adaptation mechanism to maintain dual-task performance (Al-Yahya et al., 2011). For instance, De Sanctis et al. (2014) showed that increases in task load are accompanied by increases in stride length. The authors argued that this adjustment in motor performance can be seen as a strategic mechanism to slow down walking while participants are engaged in a concurrent task. With recent advancements in wearable technology, researchers can design specialized experimental paradigms to explore the relationship between the cognitive and motor domains during increased task load.

The past decade has seen an increased interest in Mobile brain/body imaging (MoBI) studies linking brain activity to naturalistic behaviors in 3D spaces (Makeig et al., 2009; Gramann et al., 2014; Jungnickel et al., 2016). This integrative approach collects EEG, motor behavior, and environmental events to study action and participant/environment interactions (Ojeda, et al., 2014). This is critical for cognitive neuroscience given the views that human cognition has evolved to maximize behavioral success within our complex environments (Makeig et al., 2009). Furthermore, researchers have recognized that even though traditional stationary studies have advanced our knowledge about cognitive processes, that approach itself is reductionist as it takes place in artificial settings (Ladouce et al., 2017). More critically, evidence from single-cell recording studies in animals suggests that there are differences in information processing during motion as opposed to a static or resting state (Gramann et al., 2011). MoBI studies involving naturalistic behaviors such as walking and running often use high-density recordings while applying independent component analysis (ICA) to systematically separate the sources of movement artifacts from the EEG signal (Gwin et al., 2010). Having mobile flexibility in EEG research design allows for higher ecological validity than traditional laboratory studies. This opens new avenues to study cognition under naturalistic settings where participant experience matches everyday situations more than an isolated and highly controlled environment.

Previous studies from our research group have successfully recorded ERP components during an active physical performance and in real-world scenarios. For example, using an auditory oddball paradigm, Scanlon et al., (2017) reliably recorded the P3 and MMN/N2b components, as well as a frontal alpha peak, during stationary cycling. Furthermore, it has been shown that differences in the experimental environment influence the morphology of the ERP component being recorded: In a follow-up study where participants completed the oddball task while cycling outdoors, Scanlon et al., (2019) showed that the P3 and MMN/N2b are present during outside cycling. Crucially, it was found that relative to indoor cycling, there is a more negative ERP amplitude in the oddball P3 and P2 components, and a decrease of spectral power in the alpha range when cycling outside. This difference in P3 amplitude at electrode Pz was interpreted as an increase in cognitive processing due to being outdoors, where participants are continuously processing complex, external auditory and visual stimuli that compete with resources devoted to the oddball task while focusing on bicycle riding and direction.

Additionally, relative to indoors, cycling outside was associated with an increase of N1 amplitude for standard and target tones over mid frontal and parietal regions. The authors concluded that such an increase in N1 amplitude could reflect an increase in auditory filtering required to complete the oddball task in the noisier outdoor environment. These cycling studies suggest that certain ERP components show a different morphology outdoors, likely due to the influence of external stimuli bombarding the brain while being outside. Other studies comparing indoor vs outdoor cycling have also found a more negative P3 peak while participants cycle outside compared to indoors (Zink et al., 2016). In the context of dual-task interference, the decrease in P3 amplitude over parietal regions reported in these studies reflects the cognitive costs related to resource allocation during performance previously established in the literature (Nenna et al., 2020).

MoBI studies exploring CMI have also identified the P3 as one neural mechanism involved in resource allocation during task completion (Nenna et al., 2020). For example, Liebherr et al. (2018) demonstrated that increases in both motor task and cognitive difficulty lead to a more negative amplitude in the P3 time window in a cognitive task with a maximal difference over centro-parietal regions. The increase in motor difficulty was manipulated by having participants stand on one leg while completing the task. In accordance with these results, Reiser et al. (2019) employed an oddball paradigm outdoors to measure the influence of movement complexity in P3 amplitude. They showed that movement complexity was associated with increases in cognitive load followed by a more negative voltage over the P3 time window to the target tones over parietal regions. Likewise, Ladouce et al. (2019) used an oddball task to show that, relative to standing, walking is associated with a more negative P3 voltage over occipital areas to target tones. Critically, the P3 amplitude reduction in the walking condition was found to be due to visual and inertial processing during the task, and not the act of walking per se. The authors concluded that resource allocation during motion is independently modulated by inertial and visual processing. The observed modulations of P3 amplitude in these studies further show that mobile paradigms can be suitable for the study of resource allocation while in action. Taken together, these studies demonstrate that the mechanism of CMI can be successfully captured using mobile designs. In the current study, we introduce the novel electric-skateboarding (e-skateboarding; operating a self-propelling skateboard with a wireless Bluetooth handheld controller) EEG approach to test for cognitive interference in an attempt to expand the previous findings beyond the walking and cycling domains.

In addition to ERPs, oscillatory electrical brain activity has also been shown to vary with resource allocation and cognitive engagement in experimental tasks. For example, power in the alpha range (8-12 Hz) has been measured in studies of active visuospatial biasing and suppression (Kelly et al., 2009), selective inhibition, and other anticipatory mechanisms (Foxe et al., 1998; Rihs et al., 2007). Previous studies involving motor behaviors, such as walking and cycling, have found differences in alpha power based on attentional demands in these movement tasks. For example, Storzer et al., (2016) found that walking is associated with a stronger alpha decrease than stationary cycling due to an increase in sensory processing and motor planning in the walking condition. Wagner et al., (2014) have found decreases in alpha power in walking when motor planning and motor intention are required. Cycling was associated with decreases in alpha power when participants completed an oddball task riding a bicycle relative to a sitting position indoors (Scanlon et al., 2019). This decrease in power was attributed to increased stimulus processing from doing the task in an outdoor environment.

The results from the mobile EEG studies reviewed so far highlight the importance of outdoor paradigms for several reasons. First, replicating classic findings under less controlled scenarios (e.g., a busy road) is important for the field of cognitive electrophysiology since, ultimately, we want to understand and predict brain functioning in everyday situations outside laboratories. Second, it allows for exploratory and novel paradigms to be developed for research questions about complex naturalistic behaviors while maintaining higher ecological validity than stationary studies. This offers new and exciting avenues for mobile research that could validate the findings from stationary/laboratory studies (Ladouce et al., 2017). Third, mobile EEG is a promising tool that can offer affordable and flexible medical diagnosis (Krigolson et al., 2017), and can offer flexibility in measuring cognitive performance (e.g., fatigue/cognitive load) in real-life tasks (Darari et al., 2017).

In the current experiment, we used an active task of e-skateboarding on an indoor 200-meter track as a primary task while participants simultaneously completed an auditory oddball task and had their EEG recorded. We adopted a skateboard paradigm because it allows participants to be in active interaction with the environment while greatly minimizing body movements that lower the signal-to-noise ratio in EEG recordings (Oliveira et al., 2016). We employed the portable EEG methodology previously used by our research group in cycling paradigms (see Scanlon et al., 2019) where participants could freely move while wearing the EEG system in a backpack (Figure 1). Using skateboarding as the primary task, our focus was to use an auditory oddball paradigm to test whether we could record a reliable P3 and MMN effect between standard and target tones. Furthermore, we wanted to investigate whether an increase in primary task difficulty (an increase in skateboarding difficulty) would produce measurable changes in P3 amplitude. E-skateboarding difficulty can be increased by manipulating stance preference: since skateboarding requires individuals to move laterally, it is common for learners to develop a stance preference while learning the task. Preference is reflected in which foot goes forward on the board or which shoulder they look over while riding. Right-handed skateboarders generally ride with their left leg forward, however, some right-handed skaters generally prefer to ride with their right foot forward. The former riding style is generally called “regular” whereas the latter is referred to as “goofy”. Due to the stability and safety of the e-skateboard, participants can be asked to switch their stance preference in order to increase riding difficulty without turning the task into a falling hazard. In this context, riding in the non-preferred stance implies an increase in primary task difficulty, which has been previously associated with a decrease in P3 amplitude in the secondary oddball task (Kramer & Strayer, 1988; Kida et al., 2012). We tested the robustness of the oddball P3 effect while participants rode a skateboard on a running track in four different conditions (preferred stance clockwise direction, preferred stance counterclockwise direction, non-preferred stance clockwise direction, and non-preferred stance counterclockwise direction). For the ERP analysis, since dual-task processes modulate P3 amplitude, we predicted a decrease in P3 amplitude in the non-preferred stance due to increases in riding difficulty while completing the task.

**Figure 1.**
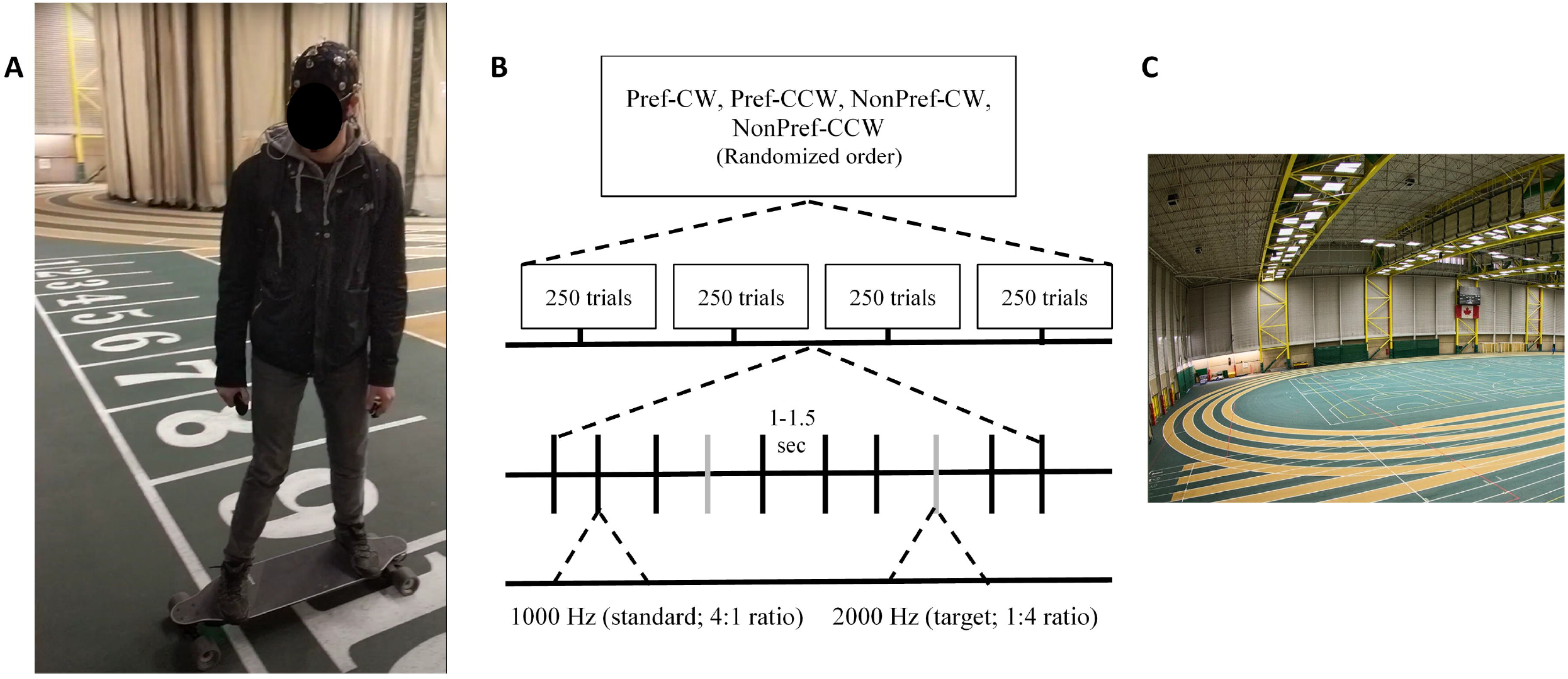
Setup task and location. A: Mobile EEG setup. B: Task diagram. C: Experiment location

Using the skateboard paradigm, we tested whether an increase in task difficulty (by having participants e-skateboard in their non-preferred stance) would be reflected in tonic alpha power decreases during the completion of the oddball task in the non-preferred condition. We also recorded participant’s resting-state EEG with their eyes both open and closed to test whether we could measure a typical alpha peak (an increase of spectra power in the 8-12 Hz range) right by the running track before completing the skateboard task. Decreases in resting-state alpha peak are associated with increased cognitive arousal (Barry, Clarke & Johnstone, 2011). Since the experiment was taking place in a busy environment, we were motivated to assess resting alpha levels prior to the start of the task. Additionally, we computed the averaged alpha power between the left and right electrodes to conduct an exploratory analysis on alpha lateralization during skateboarding. Spatial attention paradigms (Sauseng et al., 2005; Ikkai et al., 2016; Kelly et al., 2006) demonstrated that alpha power increases over areas to be ignored. Such a shift in alpha power, with enhanced power in the hemisphere contralateral to the distractors, has been identified as a mechanism of sensory inhibition (Mathewson et al., 2011).

In the current paradigm, participants ride an e-skateboard around a running track clockwise and counterclockwise. This allows us to spatially separate the source of incoming distractors based on the direction participants ride. In the clockwise condition (regardless of riding preference) the largest source of distractors (joggers, walkers, and track users) come from the right side of space while in the counterclockwise condition the source of distraction is always located on the left visual field of the participant. This is because participants were set to only ride in the outermost track during the task. We conducted exploratory analyses to test whether we could get an increase of alpha power over the parieto-occipital region contralateral to the source of distractors in the clockwise and counterclockwise riding directions.

## Methods

### Participants

A total of 29 individuals from the university community completed the study (mean age = 20.96, age-range = 18-27, goofy-footed proportion: 41.37%, right-handed proportion: 87.5% female proportion = 37.93%). Participants received either an honorarium of $20 or credits towards Research Participation in an undergraduate psychology class. All participants had normal or corrected-to-normal vision and did not have neurological antecedents. Each participant was comfortable with basic skateboarding in both preferred and non-preferred stances. This study was approved by the Internal Research Ethics Board at the University of Alberta (Pro00050069) and participants gave informed consent before completing the study. This study conforms with the World Medical Association Declaration of Helsinki.

### Materials and Procedure

The experiment was completed in the Universiade Pavilion (aka Butterdome) on a 200-meter indoor track at the University of Alberta during drop-in hours (pavilion users were performing leisure sports around the track during sessions). Participants first completed two, three-minute, resting-state tasks (beginning with either eyes closed or eyes open) to measure a baseline level of resting spectral activity while they were seated in the bleachers of the track.

Participants were instructed to breathe calmly and either keep their eyes closed or to fixate on a nearby object. Following both baseline tasks, participants were given a basic tutorial on using the Boosted V3 Stealth Electric Board (Palo Alto, California, United States; https://boostedboards.com/ca/) before starting the main skateboard experiment. The Boosted skateboard is a two-motor electric skateboard controlled by a joystick held in the right hand, with throttle and clutch controls on the thumb and trigger finger respectively. The joystick was held in the right hand while a response button was held in the left hand. Participants began the task by standing on the skateboard and pressing the response button, starting a 10-second countdown. Participants were expected to be in motion around the Universiade Pavilion track when the countdown finished. To ensure clean data, participants were instructed to remain as still as possible, blink only when necessary, and keep their gaze directed forward. A sound cue was used to instruct the participants when the task was completed. Participants rode the skateboard using a limited acceleration mode with a maximum speed of 17 kph. Participants were trained to ride at a speed at which they felt most comfortable around the track for safety purposes. On average, the total participation time was 1 hour and 50 minutes. The average riding distance was approximately 1.92 km.

Based on the participant’s preferred skateboarding stance and the direction the participant traveled around the Pavilion track, four conditions were generated: ‘preferred clockwise’, ‘preferred counterclockwise’, ‘non-preferred clockwise’, and ‘non-preferred counterclockwise’. Condition order was randomly determined and counter-balanced. In each condition, participants completed an auditory oddball task where two tones were consistently presented through headphones (either a 1000 Hz or 1500 Hz tone played at 65 dB). Participants were instructed to press the response button each time the 1500 Hz tone was played (target tone) and to withhold a response following the 1000 Hz tone (standard tone). There were no reports of issues with the task volume from any of the participants. A delay, randomly selected from a distribution between 1000 and 1500 ms, followed each tone. Response times were collected during this delay period. In all, 20% of trials contained a target tone and 80% contained a standard tone, with 250 trials in each condition. Each condition was approximately six minutes long. Between conditions, the participant returned to the starting position, the computers were reset, and the participant was given a small break.

The tones and response button task were programmed and administered using a Raspberry Pi 2 Model B computer (Raspberry Pi Foundation, Cambridge, UK) running version 9 of the Raspbian Stretch operating system and version 2.7.13 of Python. The Raspberry Pi 2 was powered via a micro-USB cable connected to a Microsoft Surface Pro 3 laptop. Audio output was via Sony earbuds connected to the 3.5mm audio connector on the Raspberry Pi 2.

Coincident in time with tone onset, 8-bit TTL pulses were sent to the EEG amplifier via a cable connected to the GPIO pins of the Raspberry Pi 2. These TTL pulses were used to mark the EEG data for ERP averaging. EEG data was collected from participants using active, wet, low-impedance electrodes (actiCAP active electrodes kept below 5 kΩ). Inter-electrode impedances were measured at the start of the experiment. The following 15 electrode locations were used, arranged in the typical 10-20 electrode positions (F3, F4, T7, T8, C3, C4, P7, P8, P3, P4, O1, O2, Fz, Cz, Pz). A ground electrode was used, positioned at AFz. Ag/AgCl disk electrodes with SuperVisc electrolyte gel and mild abrasion with a blunted syringe tip were used to lower impedances. EEG data was recorded online, referenced to an electrode attached to the left mastoid. Offline, the data was re-referenced to the arithmetically derived average of the left and right mastoid electrodes.

EEG data was recorded with a Brain Products V-Amp 16-channel amplifier (Brain Products GmbH), connected to the same Microsoft Surface Pro 3 laptop powering the Raspberry Pi, running BrainVision Recorder software (Brain Products GmbH, Gilching, Germany). In addition to the 15 EEG sensors, two reference electrodes, and the ground electrode, vertical and horizontal bipolar EOG was recorded from passive Ag/AgCl easycap disks using Bip2Aux adapters connected to the auxiliary ports of the amplifier. EOG electrodes were affixed vertically above and below the left eye and affixed horizontally 1 cm lateral from the outer canthus of each eye. The participant’s skin was cleaned using Nuprep (an exfoliating cleansing gel) (Weaver & Co, Aurora, Colorado USA) before the placement of the electrodes, electrolyte gel was used to lower the impedance of these electrodes to under 5 kΩ in the same manner as previously mentioned. Data was digitized at 1000 Hz with a resolution of 24 bits and hardware filtered online between 0.1 Hz and 30 Hz, with a time constant of 1.5155 s and a notch filter at 60 Hz.

All aforementioned equipment was held within a 2-pocket backpack worn by the participant, as shown in Figure 1. The total weight of the backpack containing the Raspberry Pi, V-vamp, and laptop was 4.55 lbs.

### EEG Analysis

Analyses were computed in MATLAB R2018a (Mathworks, Natick, Massachusetts, USA) using EEGLAB (Delorme & Makeig, 2004) and custom scripts (https://github.com/kylemath/MathewsonMatlabTools). Statistical analyses were computed on JASP (JASP Team, Amsterdam, Netherlands). The EEG markers were used to construct 1200 ms epochs (200 ms pre-stimulus baseline + 1000 ms post-stimulus) time-locked to the onset of standard and target tones, with the average voltage in the first 200 ms baseline period subtracted from the data for each electrode and trial. To remove artifacts due to amplifier blocking and other non-physiological factors, any trials with a voltage difference from baseline larger than +/-1000 µV on any channel (including eyes) were removed from further analysis. At this time, a regression-based eye-movement correction procedure was used to estimate and remove the artifact-based variance in the EEG due to blinks as well as horizontal and vertical eye movements (Gratton et al., 1983). After identifying blinks with a template-based approach, this technique computes propagation factors as regression coefficients predicting the vertical and horizontal eye channel data from the signals at each electrode. The eye channel data is then subtracted from each channel, weighted by these propagation factors, removing any variance in the EEG predicted by eye movements. Artifact rejection was again performed, using a voltage threshold of +/− 500µV. These artifact rejection thresholds were chosen to be relatively lenient, similar to other mobile EEG studies we have done (see Scanlon et al., 2017; Scanlon et al., 2019; Scanlon et al., 2020), to quantify how much noise (electrical, muscle, or otherwise) was present in the ERP data and to ensure an adequate number of trials were available for analysis. Baseline correction was performed again following the second artifact rejection. Table 1 shows the mean trial count for target and standard tones used for each condition after artifact rejection. The rejected number of trials for targets and standards respectively was similar for all conditions(*F*(3,112) = 0.84, *p* = 0.48).

**Table 1:** Trial count information for targets and standards in each condition.

| Condition | Stimulus Type | Mean Trial Count | Standard Deviation | Trial Count Range (Minimum:Maximum) |
| --- | --- | --- | --- | --- |
| Preferred Clockwise | Targets | 46.90 | 4.87 | 29:50 |
|  | Standards | 185.83 | 21.61 | 104:200 |
| Preferred Counterclockwise | Targets | 47.79 | 5.06 | 26:50 |
|  | Standards | 191.52 | 17.35 | 115:200 |
| Non-preferred Clockwise | Targets | 47.90 | 4.47 | 30:50 |
|  | Standards | 190.34 | 17.59 | 131:200 |
| Non-preferred Counterclockwise | Targets | 47.76 | 4.56 | 29:50 |
|  | Standards | 189.79 | 20.75 | 92:200 |

### ERP Analysis

Appropriate time window cut-offs for the MMN/N2b and P3 waveforms were determined by creating grand-average ERP waveforms, averaged across all participants and conditions, to create a single ERP waveform for electrodes Fz and Pz (Figure 2). We selected the peak within the grand average waveform as the center of the component of interest and used a window of 150 ms for the P3 analysis and a window of 100 ms for the MMN/N2b. These grand-average waveforms were used to avoid biasing the selected time windows towards any one condition.

**Figure 2.**
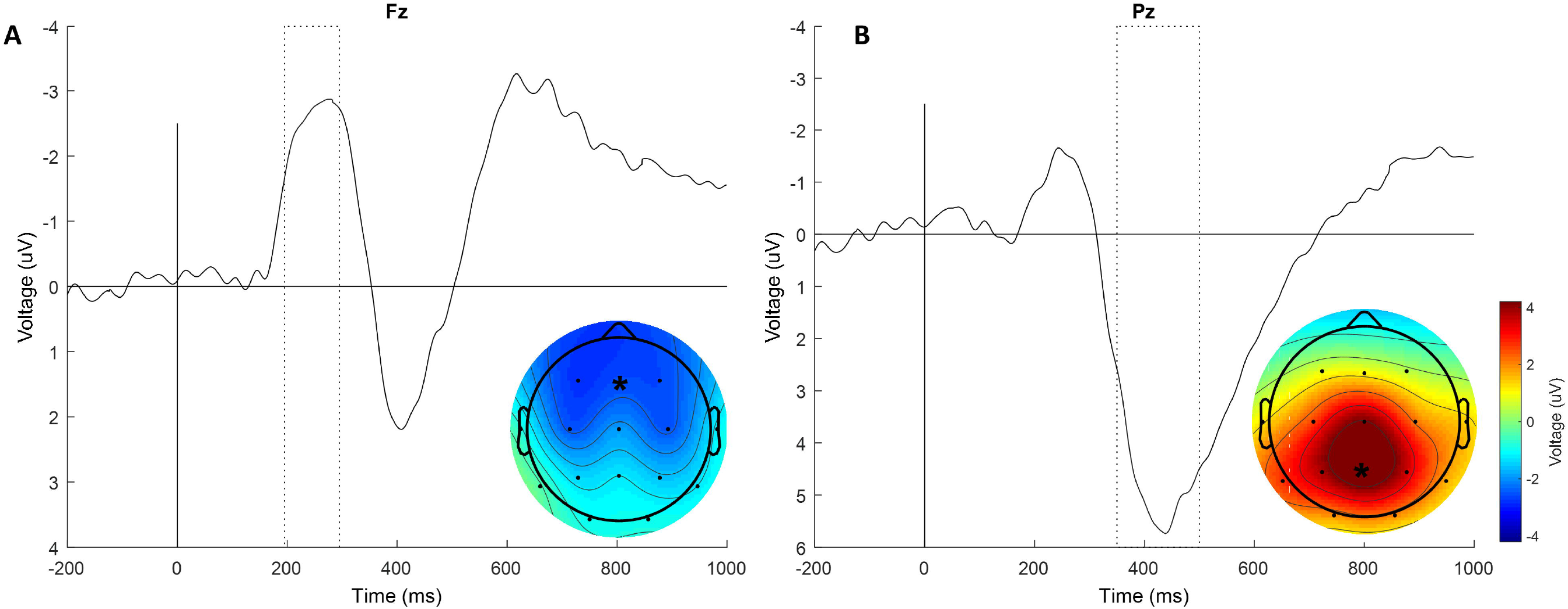
Grand-averaged ERP across conditions used to select time windows for analyses.

The negative deflection between 195-295 ms for electrode Fz was used for the MMN/N2b time window while the large positive deflection between 375-525 ms for electrode Pz was used for the P3 time window. Trial-averaged ERPs were derived in each of the four conditions (‘preferred clockwise’, ‘preferred counterclockwise’, ‘non-preferred clockwise’, and ‘non-preferred counterclockwise’) and waveforms were compared. A 2×2×2 ANOVA was run to compare tone type (standard and target), stance preference (preferred and non-preferred), and clock orientation (clockwise and counterclockwise) during the MMN/N2b and P3 time windows, at electrodes Fz and Pz respectively. Two-way repeated-measures ANOVAs were also conducted on the difference waveforms for the MMN/N2b and P3 time windows at electrode Fz and Pz respectively. Stance preference and clock orientation were the factors used for these analyses.

These waveforms were derived by subtracting the standard ERP waveforms from the target ERP waveforms for both the preferred/non-preferred and clockwise-counterclockwise conditions, with α set to 0.05 for all analyses.

### Spectral Analysis

Average frequency spectra across the baseline and oddball epochs was also computed using the wavelet routine from the Better Oscillation Method (BOSC; Whitten et al., 2011) with a 6-cycle wavelet transform across a frequency range of 0.1 Hz to 30 Hz, increasing in 0.5 Hz steps. We chose this wavelet cycle parameter because we were interested in the overall sustained alpha power instead of the differences in power between target and standard tones. Spectral data was further analyzed using the “Fitting Oscillations & One Over F” (FOOOF) algorithm (Haller et al., 2018) to represent the data as two distinct components; the aperiodic background 1/*f* and periodic oscillations which may contain greater spectral power than the aperiodic component.

This analysis was performed using version 0.1.1 of the FOOOF Matlab wrapper with the following settings: peak width limits = [0.5,12]; max number of peaks = Inf; minimum peak amplitude = 0.0; peak threshold = 2.0; background mode = fixed; verbose = true; frequency range = 0.5-30 Hz. The background 1/*f* spectra was then subtracted from the periodic component to better compare changes in spectral power between 0.5 Hz and 30 Hz across our conditions. All spectral analyses were done using this calculated FOOOF spectral data (Figure 6 C-D).

For the baseline task, spectra were computed for each participant by first dividing the data into 3000 ms segments. Spectra were calculated for each chunk, which was then averaged across the 3000 ms duration for each chunk. Then each averaged-chunk was combined within each participant and finally averaged across participants to generate grand-averaged spectra for both eyes open and eyes closed conditions. For the oddball task, we generated 3000 ms epochs around the onset of each standard trial (1000 ms pre and 2000 ms post standard onset) for each participant. Spectra were calculated for each epoch, then averaged across time and number of standard trials to generate spectra for each participant. These spectra were then averaged across participants to create grand-average spectra for each condition in the oddball task. Spectra were calculated using electrodes Fz and Pz. Resting-state EEG was not recorded from two participants and therefore their data was excluded from this main spectral analysis. Furthermore, to explore the role of left-right visual field distractors in hemispheric alpha, power was computed by combining the following parieto-occipital electrodes: left hemisphere (P7, P3, O1), right hemisphere (P8, P4, O2).

## Results

### Behavioral Results

Figure 3A shows the mean accuracy in response to target tones across all four task conditions and by global stance preference (collapsed over direction of travel). Results from a two-way repeated measures ANOVA on behavioral accuracy show no significant main effect for either preference, (*F*(1, 28) = 0.74, *p* = 0.40, η^2^_p_ = 0.03), or clock orientation (F(1, 28) = 0.85, p= 0.36, η^2^_p_ = 0.03) and no significant interaction (*F*(1, 28) = 0.01, *p* = 0.91, η^2^_p_ = 4.61e-4).

**Figure 3.**
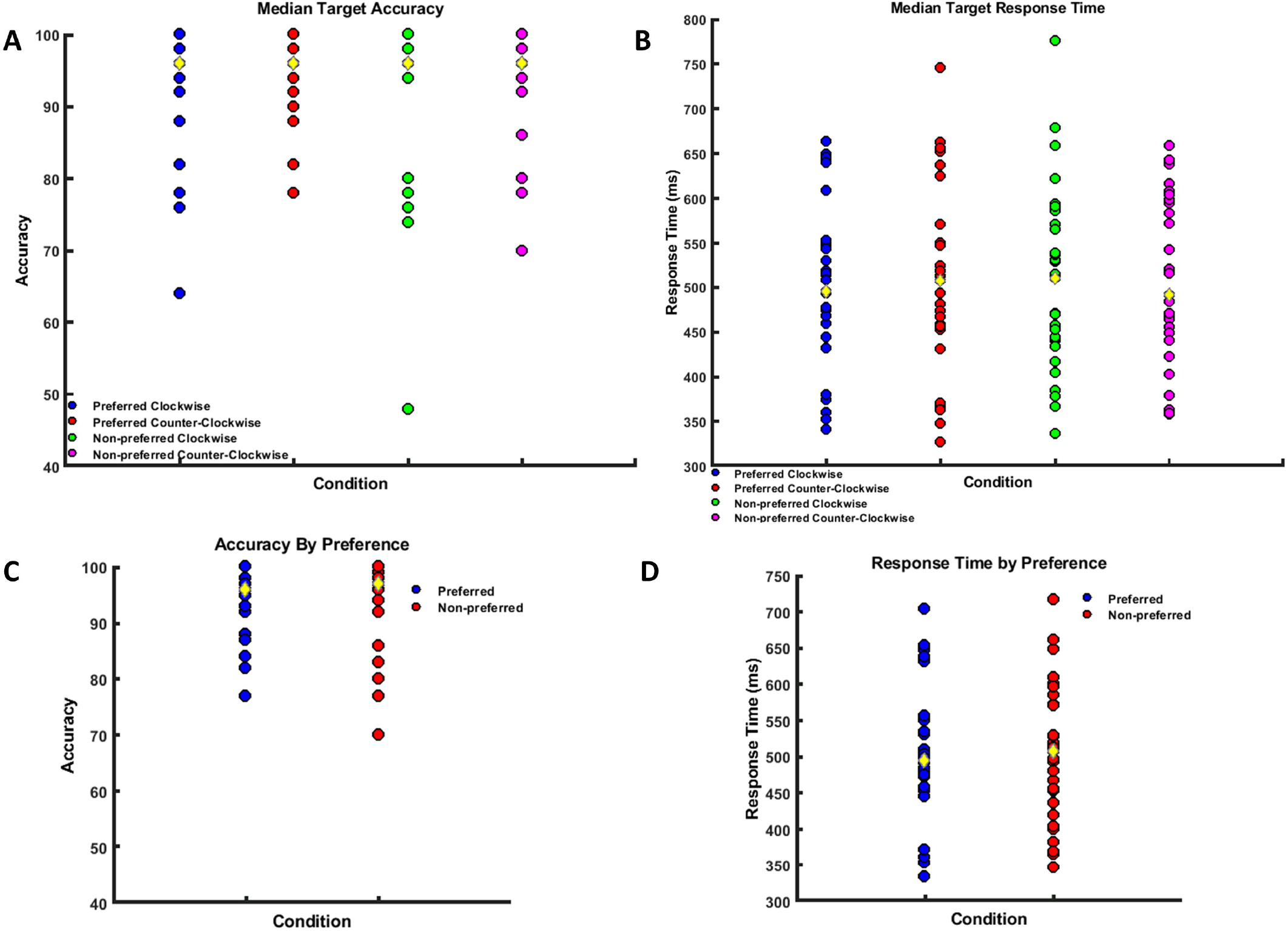
Behavioral task results. A: Mean target accuracy. B: Median target reaction time. C: Target accuracy by grand preference. Target reaction time by grand preference

Figure 3B shows mean reaction times in milliseconds for all four task conditions and by global stance preference. A two-way repeated measures ANOVA on reaction times show no significant main effect for either preference, (*F*(1, 28) = 2.56e-4, *p* = .99, η^2^_p_ = 9.15e-6), or clock orientation (*F*(1, 28) = .03, *p* = 0.87, η^2^_p_ = 9.29e-4), and no significant interaction (*F*(1, 28) = 0.27, *p* = 0.61, η^2^_p_ = 0.01).

### ERP Morphology and Topography

Figure 4A shows the grand average ERPs for target and standard tones at electrode Fz for all study conditions. Target tones are depicted in colored lines while standard tones are depicted in black lines. Standard errors at each time point are depicted by the shaded regions. Compared to standard tones, there is an increase in negative voltage for the MMN/N2 time window between 195 and 295 ms following target tones. Figure 4C shows the grand average ERP at electrode Pz for all conditions, showing a clear positive increase in voltage in the P3 window (350 and 550 ms) following target tone onset. Additionally, topographies of the target-standard difference were computed for the MMN/N2b and P3 time windows across conditions to show the overall scalp activation distribution for both time windows. Based on the time window chosen for these topographies, one can observe a more anterior distribution for the MMN/N2b time window and a more posterior distribution for the P3 time window. We conducted a 2×2×2 repeated measures ANOVA to understand the relationship between target/standard tones in all four conditions and whether the MMN/N2b and P3 are influenced by riding condition. For the MMN/N2b time window at electrode Fz, there was a significant main effect for tone type (*F*(1, 28) = 87.72, p < .001, η^2^_p_ = 0.76). There was no significant effect of preference type (*F*(1, 28) = 0.63, p = 0.43, η^2^_p_ = 0.02) or orientation (*F*(1, 28) = 2.78, p = 0.11, η^2^_p_ = 0.09), and there were no significant interactions: tone type∗preference (*F*(1, 28) = 2.04, p = 0.17, η^2^_p_ = 0.07), tone type∗orientation (*F*(1, 28) = 0.92, p = 0.35, η^2^_p_ = 0.03), preference∗orientation (*F*(1, 28) = 1.87, p = 0.18, η^2^_p_ = 0.06), tone type∗preference∗orientation (*F*(1, 28) = 0.01, p = 0.91, η^2^_p_ = 4.77e-4). Table 2 shows the significant pairwise comparisons where it can be seen that the target-standard difference at the MMN/N2b window was reliable in all four conditions. All p-values for the pairwise comparisons were adjusted for multiple comparisons using Bonferroni correction.

**Table 2:** Pairwise results comparing target and standard ERP waveforms in each condition.

| Comparison | Electrode/ Time Window | Mean Voltages (μV)<br>(Targets:Standards) | $t(28)$ | $p$ | Confidence Interval | Effect Size (Cohen's $d$) |
| --- | --- | --- | --- | --- | --- | --- |
| Preferred Clockwise | Fz/MMN-N2b | -3.23:-0.58 | 5.43 | <0.001 | -4.23:-1.09 | -1.23 |
|  | Pz/P3 | 4.71:-0.47 | 5.98 | <0.001 | 2.36:8.01 | 1.32 |
| Preferred Counterclockwise | Fz/MMN-N2b | -2.86:-0.72 | -4.37 | <0.001 | -3.71:-0.57 | -0.90 |
|  | Pz/P3 | 4.35:-0.54 | 5.64 | <0.001 | 2.07:7.71 | 0.94 |
| Non-preferred Clockwise | Fz/MMN-N2b | -3.96:-0.75 | -6.56 | <0.001 | -4.78:-1.64 | -0.97 |
|  | Pz/P3 | 4.59:-0.78 | 6.20 | <0.001 | 2.55:8.20 | 1.18 |
| Non-preferred Counterclockwise | Fz/MMN-N2b | -3.10:-0.30 | -5.71 | <0.001 | -4.37:-1.23 | -1.10 |
|  | Pz/P3 | 3.95:-0.05 | 4.61 | <0.001 | 1.17:6.82 | 0.82 |

**Figure 4:**
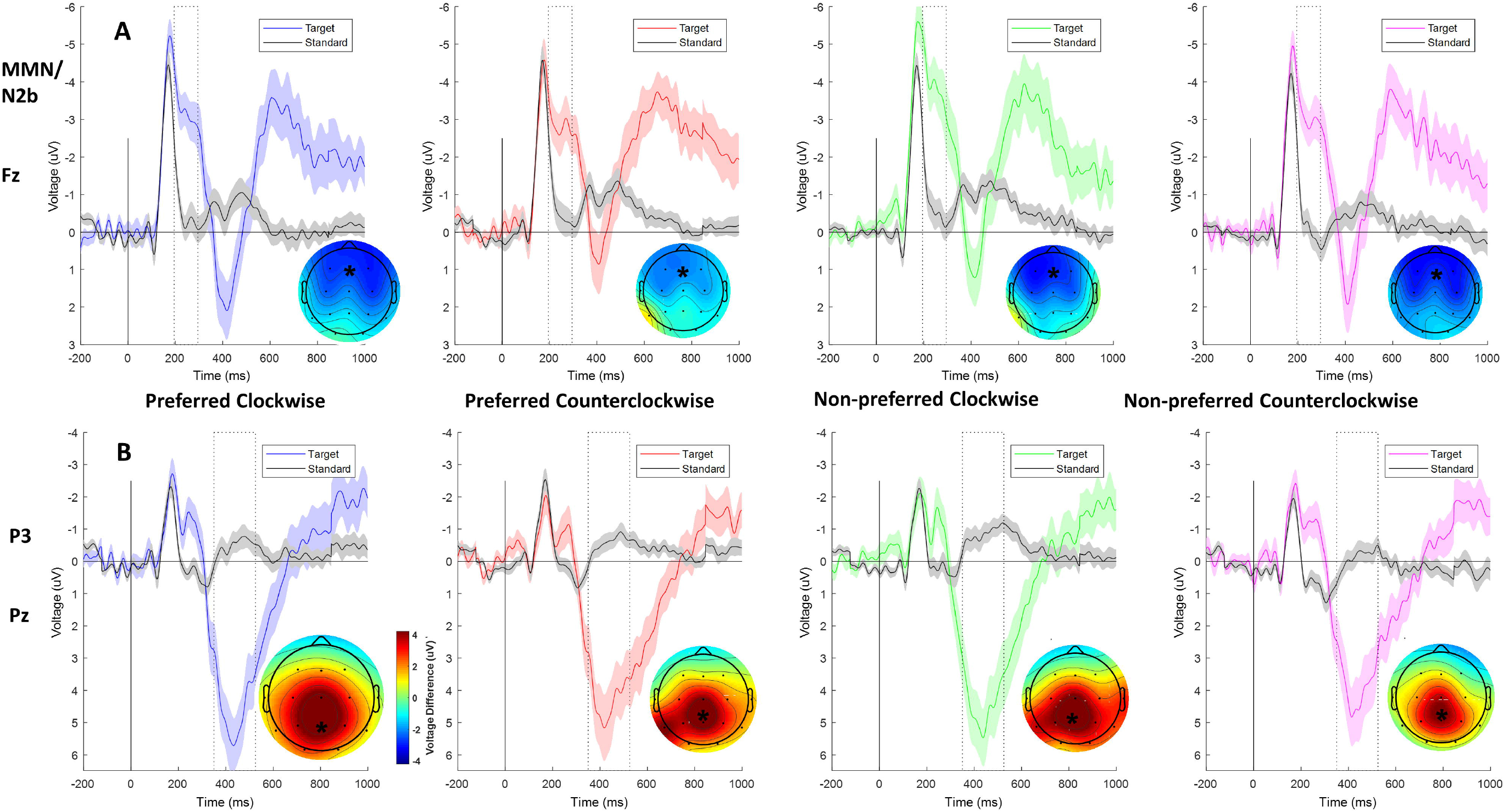
A: ERPs at electrode Fz. Conditions (left to right): Preferred clockwise, preferred counterclockwise, non-preferred clockwise, non-preferred counterclockwise. Target tones: colored lines. Standard tones: black lines. Shaded regions represent standard errors. Topographies for MMN/N2 window. B: ERPs at Electrode Pz. and topographies for P3 time windows.

For the P3 window at electrode Pz, there was a significant main effect for tone type (*F*(1, 28) = 50.86, *p* < .001, η^2^_p_ = 0.65), no significant effect of preference type (*F*(1, 28) = 0.08, p = 0.78, η^2^_p_ = 3e-3) or orientation (*F*(1, 28) = 0.11, p = 0.74, η^2^_p_ = 4e-3), and there were no significant interactions: tone type∗preference (*F*(1, 28) = 0.41, p = 0.53, η^2^_p_ = 0.01), tone type∗orientation (*F*(1, 28) = 1.42, p = 0.24, η^2^_p_ = 0.05), preference∗orientation (*F*(1, 28) = 0.17, p = 0.68, η^2^_p_ = 6e-3), tone type∗preference∗orientation (*F*(1, 28) = 0.83, p = 0.37, η^2^_p_ = 0.03). Table 2 shows the pairwise comparisons at electrode Pz where, across all conditions, the oddball P3 difference between targets and standards was highly reliable across all combinations of orientation and preference.

We also analyzed the difference waveforms of our ERPs across the preference and orientation conditions. These waveforms were calculated by subtracting standard trial ERPs from target trial ERPs, allowing us to better understand the differences in evoked activity between our two-tone types. Figure 5A shows the difference-wave ERP for the preferred and non-preferred stance in the MMN/N2b time window. Figure 5B shows the difference-wave ERP for the clockwise and counterclockwise conditions in the MMN/N2b time window. A 2×2 repeated measures ANOVA was calculated for electrodes Fz and Pz and their respective time windows, as described previously. Our results at electrode Fz show no significant effect of preference (*F*(1, 28) = 2.04, p = 0.17, η^2^_p_ = 0.07) or orientation (*F*(1, 28) = 0.92, p = 0.35, η^2^_p_ = 0.03), and no significant interaction (*F*(1, 28) = 0.01, p = 0.91, η^2^_p_ = 4.78e-4).

**Figure 5.**
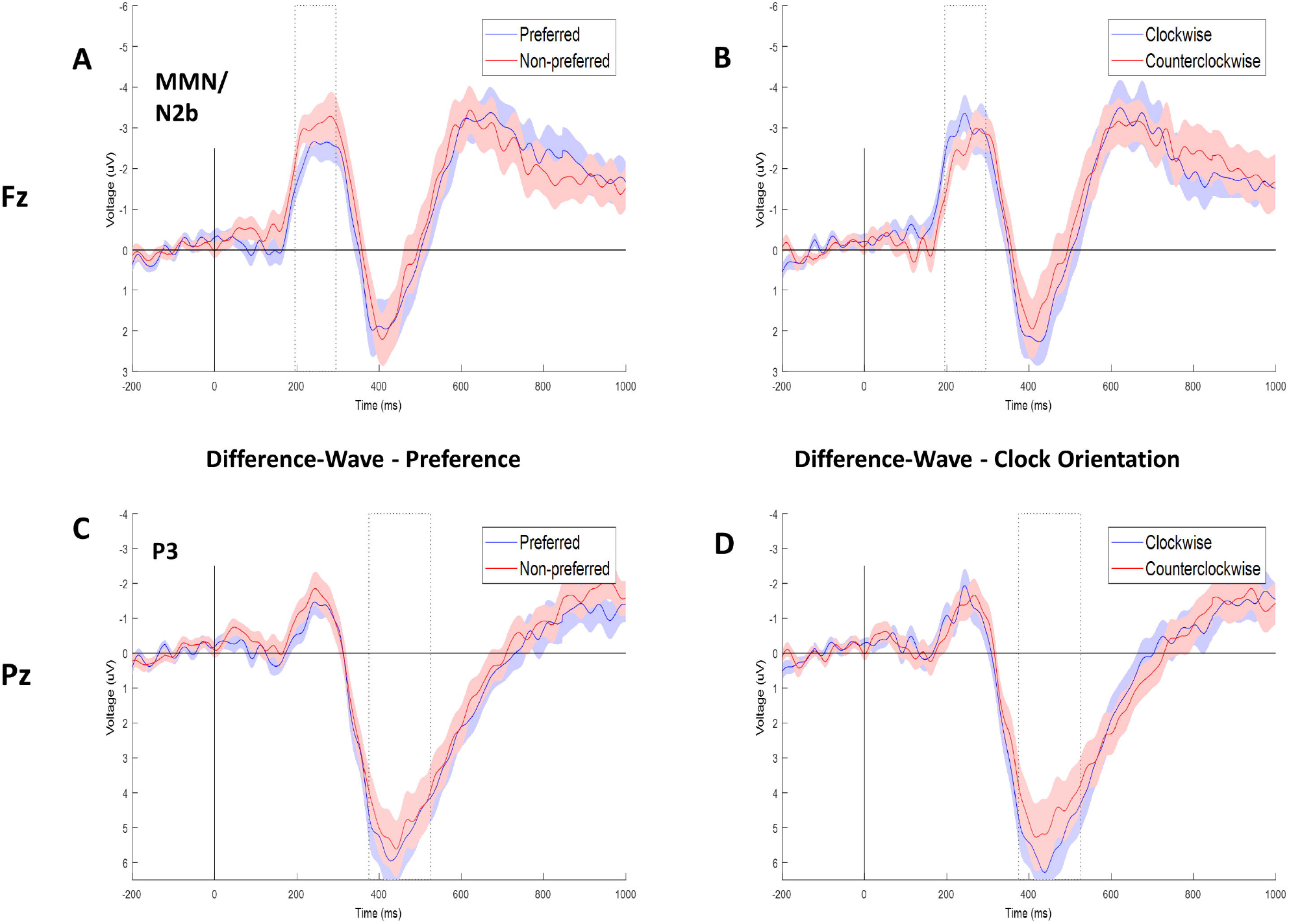
Difference-wave ERPs by A: grand preference for the MMN/N2b time window at electrode Fz. B: grand clock orientation for the MMN/N2b time window at electrode Fz. C: grand preference for the P3 time window at electrode Pz. D: grand clock orientation for the P3 time window at electrode Pz.

**Figure 6.**
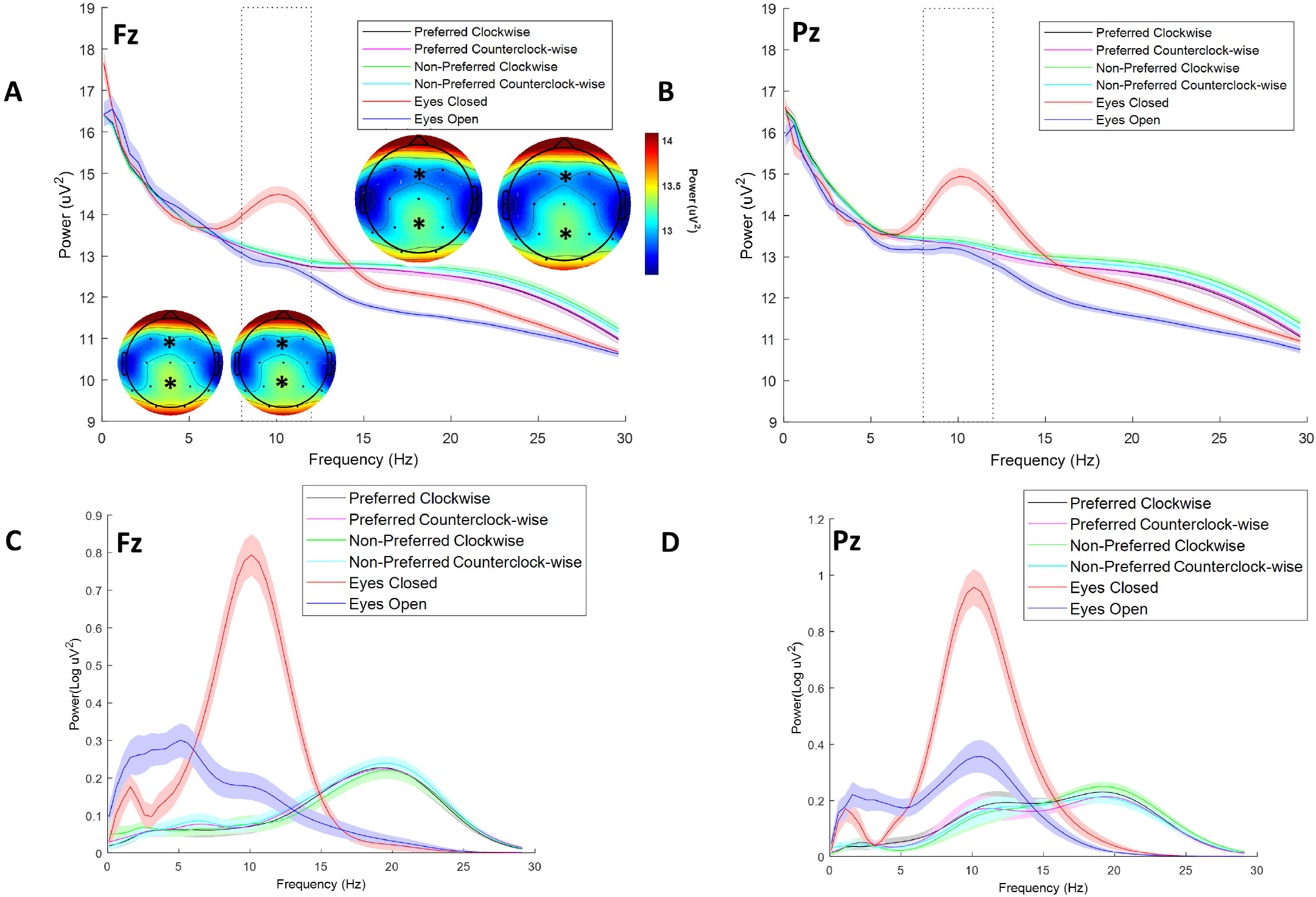
A: Power spectra at electrode Fz for all conditions plus resting-state baseline (eyes opened/closed). Shaded regions represent standard errors for all conditions. Power topographies for alpha band range (8-12Hz) for preferred clockwise and counterclockwise conditions (upper region) and non-preferred clockwise and counterclockwise conditions (lower region). B: Power spectra at electrode Pz for all conditions. C: FOOOF-background data used for statistical analysis at electrode Fz. D: FOOOF-background data used for statistical analysis at electrode Pz.

Figure 5C shows the difference-wave ERP by preference conditions at electrode Pz for the P3 time window. Figure 5D shows the difference-wave ERP for the clockwise and counterclockwise conditions in the P3 time window. Similar results also occur for electrode Pz, with no significant effect of preference (*F*(1, 28) = 0.41, p = 0.53, η^2^_p_ = 0.02), or orientation (*F*(1, 28) = 1.42, p = 0.24, η^2^_p_ = 0.05), and no significant interaction (*F*(1, 28) = 0.83, p = 0.37, η^2^_p_ = 0.03).

### Spectral Results

Figure 6A shows the power spectra plots for all four conditions for the frequency range 1-30 Hz at electrode Fz. Resting-state baseline spectra for eyes open-close conditions is also depicted. These resting-state power spectra show a predominant peak within the alpha frequency range (8-12 Hz) which is larger during the eyes-closed condition. Power topographies for the preferred and non-preferred clockwise conditions (upper region) and preferred and non-preferred counterclockwise conditions (lower region) were generated for the alpha frequency range.

Topographies show higher power distribution across central parietal-occipital regions, particularly in the non-preferred conditions. Figure 6B shows the power spectra for all conditions and baseline at electrode Pz. Figures 6C-D depict the FOOOF-background EEG spectra used for statistical analysis. This spectra plot shows a clear peak in the alpha frequency in the resting state condition that is more predominant in the eyes-closed condition. These spectra plots also reveal an increase in the Beta activity (13-30 Hz) in all the riding conditions relative to the eyes open spectra at electrodes Fz and Pz likely due to muscle activity. Relative to the resting state, there is a general decrease of alpha power in the riding conditions. A one-way repeated measures ANOVA was conducted to assess the differences in alpha peak for all conditions at electrodes Fz and Pz. A Greenhouse-Geisser correction of sphericity was applied to the analysis (ε_Fz_ = 0.38, ε_P z_ = 0.44). Significant effects of condition type were observed for electrode Fz (*F*(1.91, 49.66) = 84.84, p < 0.001, η^2^_p_ = 0.76) and electrode Pz (*F*(2.12, 57.50) = 58.01, p < 0.001, η^2^_p_ = 0.69).

Table 3 shows the significant pairwise comparisons. All p-values were adjusted for multiple comparisons using Bonferroni correction. This analysis shows that both resting-state conditions (eyes open and eyes closed) are significantly different from all skateboard conditions at electrode Pz and marginally significant at electrode Fz. There are no further significant differences between conditions. We conducted a two-way repeated-measures ANOVA using preference and clock orientation as factors at electrodes Fz and Pz. We did not find a main effect for either riding preference (F(1, 26) = 0.38, p = 0.57, shows the significant = 0.01) or riding orientation (*F*(1, 26) = 3.46, p = 0.07, η^2^_p_ = 0.009). No significant main effects were found at electrode Pz for preference (*F*(1, 26) = 1.65, p = 0.20, η^2^_p_ = 0.060) or riding orientation (*F*(1, 26) = 0.2, p = 0.603, η^2^_p_ = 0.01).

**Table 3:**
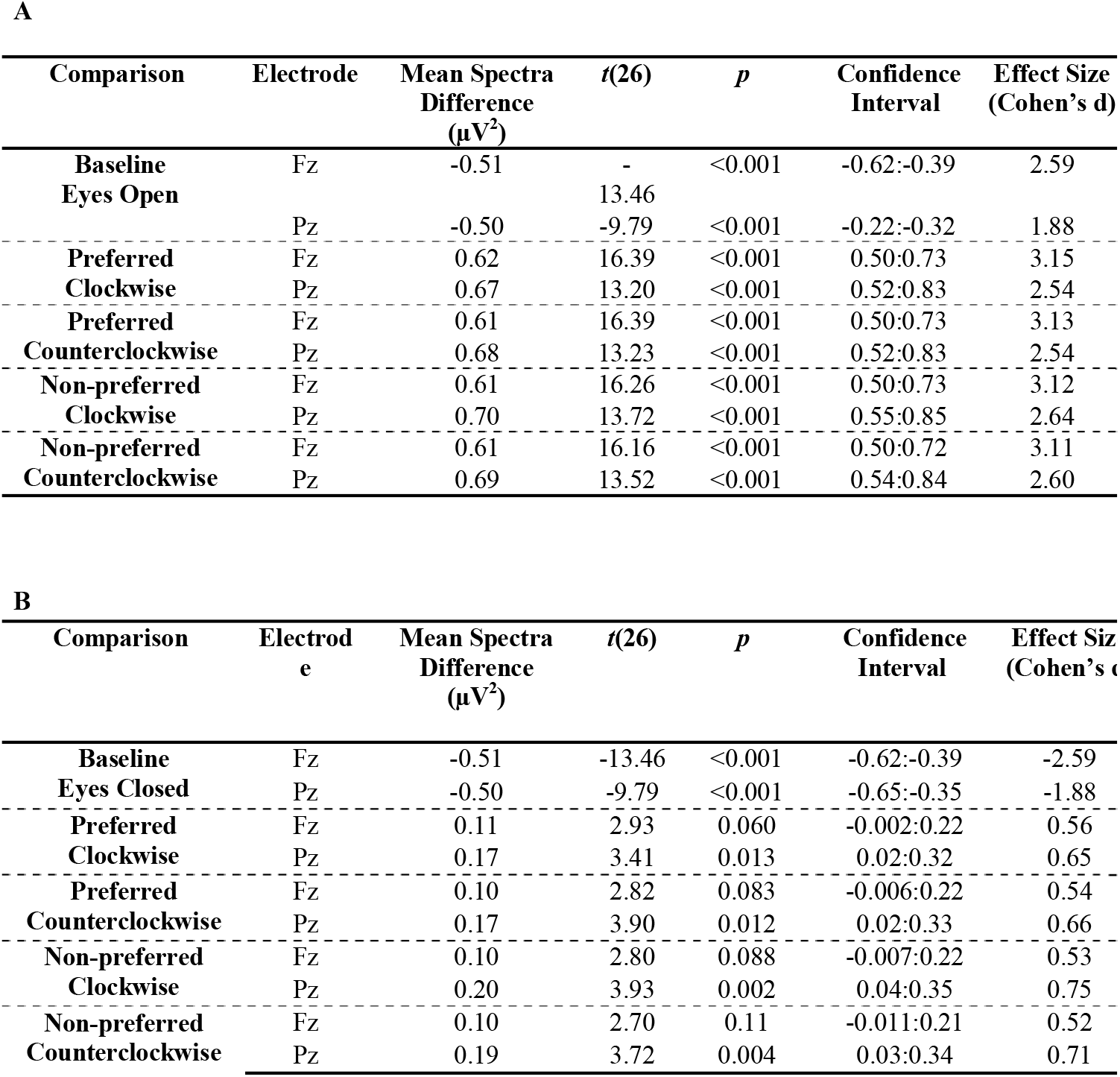
Pairwise results comparing A) Baseline Eyes Closed and B) Baseline Eyes Open spectra to each of the other five conditions.

### Left-Right Hemispheric Alpha by Preference and Riding Orientation

We conducted an additional exploratory analysis to measure whether peripheral distractors influence alpha lateralization (due to the role of alpha synchronization in suppressing irrelevant areas of the visual field), we compared the left minus right hemisphere difference in alpha band power based on riding orientation (clockwise vs counterclockwise).

Figure 7 shows the left and right parieto-occipital alpha power by riding direction in the clockwise (left side panel) and counterclockwise (right panel) conditions. We conducted a t-test of the power difference between left-right hemispheres in the clockwise and counterclockwise conditions. Such t-test revealed no differences in hemispheric alpha power between conditions (*t*(28) = 0.12, *p* = .90 d = 0.02).

**Figure 7.**
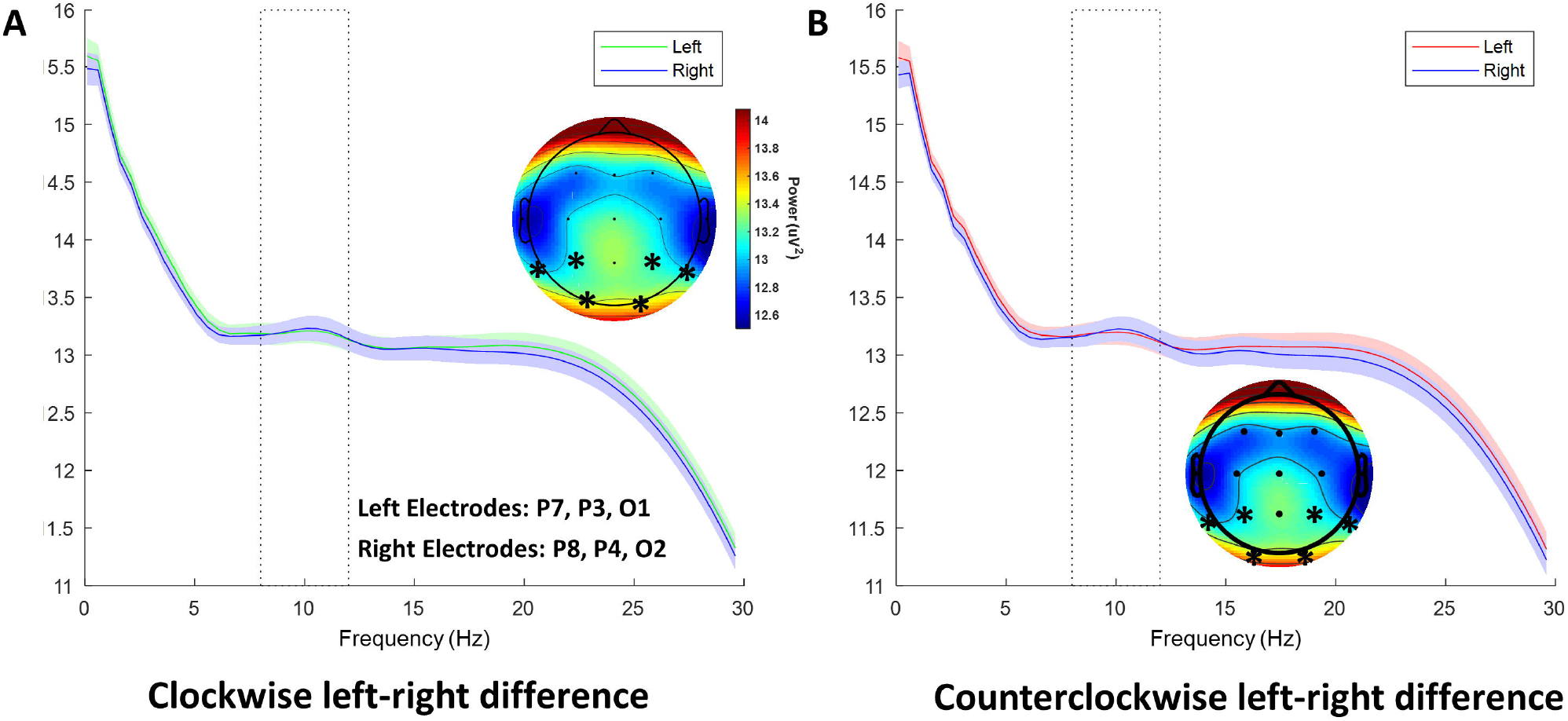
Power spectra for left and right parieto-occipital hemispheres for clockwise direction (left) and counterclockwise direction (right).

## Discussion

The present study employed a mobile EEG paradigm to successfully record ERP components and oscillatory activity from participants moving on an e-skateboard. Participants responded to an auditory oddball task while riding around a complex scenario in their preferred and non-preferred stances. We introduced a higher degree of motor interference in the non-preferred condition to evaluate whether there was a change in resource allocation measured by the posterior P3 amplitude. To our knowledge, this is the first portable EEG experiment using skateboarding. We chose to use a skateboarding paradigm for several reasons: We wanted to deploy the mobile EEG methodology previously developed by our research group (Kuziek et al, 2017; Scanlon et al, 2017) to a novel scenario using a new task, as it is important to demonstrate that this mobile EEG methodology can help researchers design and test attentional paradigms away from the laboratory. Our study further demonstrates the robustness of the oddball-P3 effect under a novel skateboard task, providing further evidence that classic attentional paradigms can be replicated under less controlled environments and complementing a long history of laboratory research using this task (Kok, 2001). The implementation of the e-skateboards (and e-scooters) offers a simple solution to the problem of excessive motion artifacts during EEG data collection while allowing participants to freely move in space. Considerable efforts have been made to ensure that the contribution of motion artifacts in the EEG data is managed effectively, such as the integration of portable EEG in experiments involving movement and action. For example, the MoBI approach (see Makeig et al., 2009; Gramann et al., 2010; Gwin et al,.2010) simultaneously records EEG, muscle movements, and environmental events to study cognitive processes away from the traditional EEG approach that requires substantial restrictions in human behavior. The groundbreaking MoBI approach has adopted multi-data streaming that includes electromyographic sensors and relies on advanced decomposition techniques to achieve the ultimate goal of exploring human cognition under natural and unconstrained behavior. According to Gramann (2014) “new techniques are now required for studying cognition under a more general range of conditions that include natural motor behavior” (p. 1). The present skateboarding approach is a simple but highly effective tool that can help mobile researchers accomplish that goal of conducting research while subjects can perform unconstrained navigation. As it was pointed out by Nenna and colleagues (2020), many MoBI studies have used artificial setups (e.g., treadmills in walking studies) where motor behaviors are not captured as they would occur in everyday fashion. We deployed the skateboard paradigm to a running track but future studies can make use of different environments (parks, roads, etc.) to achieve great ecological validity while exploring cognition. Given that balance during e-skateboarding can be easily accomplished by maintaining a proper center of gravity (e.g., keeping one’s legs spread well enough), this new paradigm can be a great tool for mobile EEG researchers who want to adopt a naturalistic task that does not introduce excessive motion artifacts during recording.

### Behavioral Results

In the current study, participants responded to target tones by pressing a button with their left hand. We calculated response accuracy and reaction time based on the four experimental conditions (preferred-clockwise, preferred-counterclockwise, non-preferred-clockwise, and non-preferred-counterclockwise) and by the grand averaged preferred and non-preferred conditions. We found no significant differences in accuracy or reaction time across conditions, even though accuracy was marginally higher in the preferred stance condition. Additionally, there was a larger standard deviation in the non-preferred condition for both accuracy and reaction time, suggesting that participants performed more uniformly in the preferred stance. These results contrast previous findings, such as Reiser et al. (2019), where increases in motor task difficulty in an oddball paradigm were associated with poorer task performance. It is possible that behavioral effects could not be achieved by our oddball task due to a ceiling effect since mean accuracy was quite high in all conditions.

### P3 & MMN/N2b Results

We hypothesized that the target-related P3 would be larger in the preferred stance condition due to differences in riding difficulty. Previous research shows that an increase in task difficulty (i.e., riding in one’s least comfortable stance) should be reflected in a more negative amplitude voltage in the P3 time window (Kok, 2001; De Sanctis et al., 2014). While we were able to measure significant, and expected, differences between ERPs following standard and target tones (we were able to detect a P3 response following rare stimuli as shown in Figure 2B and Table 2) we found no significant P3 differences between our conditions. Previous mobile EEG studies in naturalistic environments have shown a P3 amplitude decrease during increased task load in walking (Ladouce et al., 2019) and cycling (Zink et al., 2016; Scanlon et al., 2019; Scanlon et al., 2020). We expected that the increase in difficulty in the non-preferred riding condition was going to be marked by a more negative voltage within the P3 time window.

However, we failed to find differences in our current paradigm. Similar results were reported by Gramann et al. (2010), where they show no differences in P3 amplitude based on increases in motor demands in a visual oddball task. Interestingly, previous findings from Ladouce and colleagues (2019) show that the P3 amplitude reduction seen during cognitive interference is not modulated by motor demands from the act of walking itself. These results go against the view that increases in task load in the motor domain lead to a reduction of cognitive resources (Leone et al., 2017). Additionally, Ladouce has shown that resource allocation during movement is modulated by both inertial and visual stimulation at the sensory level. It could be possible the similarity in visual and inertial stimulation during the preferred and non-preferred conditions might have modulated the parietal P3 amplitude in the same magnitude regardless of riding difficulty. If this is the case, one can expect a similar level of reduction in P3 amplitude during e-skateboarding regardless of a possible increase in riding difficulty. Considering that no behavioral differences in accuracy and reaction time were found by increasing riding difficulty, it is possible that the change in stance preference did not interfere with the cognitive resources employed by the oddball task. Future paradigms should consider increasing the oddball task difficulty (e.g., manipulating tone frequency), or the non-preferred task difficulty (e.g. wobbly skateboards).

We also found an MMN/N2b difference between target-standard tones in all our experimental conditions. These results replicate previous EEG findings from other cycling studies by our research group (Scanlon et al., 2017; Scanlon et al., 2019). This ERP component is elicited by sudden changes or deviations following the repeated presentation of a stimulus, such as the change from target to standard tones (Patel & Azzam, 2005). While the standard-target amplitude difference is present in all our conditions, we did not find differences in this component based on riding preference. The lack of differences in MMN/N2b amplitude is likely due to the environmental noise being the same in all task conditions for participants, reflected in similar ERP amplitudes across conditions.

### Spectral Results

We recorded each participant’s eyes open/close baseline spectra before completing the oddball task. We found an expected and significant increase in power within the alpha range (8-12 Hz) at electrodes Fz and Pz for the eyes-closed condition compared to eyes-open, as shown in Table 3. Additionally, we found a complete attenuation of alpha power during the skateboard task relative to both baseline conditions at electrode Pz. At electrode Fz, this general reduction of alpha power between resting state and riding was found to be overall marginally significant at electrode Fz (table 3). These results support the findings of previous studies showing a decrease in alpha during complex behaviors such as walking (Beurskens et al., 2016; Storzer et al., 2016) and bicycle riding (Zink et al., 2016; Scanlon et al., 2020). However, it is important to note that this reduction in alpha power should not be taken as evidence that the experimental manipulation leads to quantifiable reductions in alpha since participants did not complete the oddball task during the resting conditions. One of the goals of the present study was to assess whether increases in riding difficulty would be reflected in a reduction in the alpha band, based on our understanding that alpha power is modulated by increases in task demand (Foxe, et al., 2011).

However, there were no significant differences in alpha power based on riding preference. Since we found an overall decrease in power during the task relative to their resting state power, it is possible that the desynchronization in alpha power we found is likely due to an overload of incoming stimuli from the task and environment that would induce a general state of cortical excitability (Klimesch et al., 2011). We found an unexpected increase of power in the beta band (13-30 Hz) in the skateboarding conditions relative to the resting spectra (Figure 6). We attribute this increase of power to increased muscle movements. Similar results were reported by Scanlon et al., (2019), showing an overall increase in beta power during outdoor cycling. This beta increase during the riding conditions contradicts previous findings linking beta desynchronization to active movements (Jain et al., 2013).

An exploratory goal in the present study was to measure the potential influence of peripheral distractors in alpha lateralization suppression. Generally speaking, participants consistently experienced a higher number of distractors in the inner tracks of the running pavilion due to joggers and track users present while participants completed the experimental task. Based on previous findings showing shifts in hemispheric alpha based on the location of spatial distractors (Sauseng et al., 2005; Kelly et al, 2006; Ikkai et al., 2016), we were motivated to compare parieto-occipital alpha power across hemispheres to measure whether we could get an effect in overall lateralization without systematically manipulating distractor location. We found no significant differences in mean alpha power in parieto-occipital regions. In a study by Malcolm et al., (2018) exposure to optic flow was found to attenuate parietal alpha power during walking. It might be possible that the state of motion experienced by the skateboarders led to a consistent suppression of induced alpha power between conditions. We did not find any reliable effects in terms of alpha lateralization, but more suitable cueing paradigms that fully isolate distractor location could help in answering whether alpha lateralization to visual distractors during active motion occurs.

### ERSP Plans

We should acknowledge that we did not look at task-evoked activity in this, or in any previous study from our research group. This is an important avenue for future mobile studies interested in exploring short term changes in cortical excitability evoked by the task events.

Changes in evoked power within certain frequencies (e.g., decreases in alpha/increases in beta) following periods of interest reflect the allocation of attentional resources that facilitate task performance (Mathewson et al., 2011). For instance, authors have previously shown event-related desynchronizations in the alpha band during the P3 period in an oddball task (Bernat et al., 2007). Since desynchronization in alpha power has been associated with increases in excitability and information processing (Klimesch et al., 2007), it would be relevant to explore evoked changes in alpha power during motion in the current paradigm. Additionally, increases in frontal theta power have been previously associated with increases in cognitive load (Di Flumeri et al., 2018), and during periods of response inhibition thereby reflecting increases in executive control (Nigbur et al., 2011). Exploring the temporal dynamics of frontal theta activity during preferred and non-preferred riding could be an interesting avenue to pursue since changes in mean theta activity have been linked to levels of mental effort devoted to task completion (Onton et al., 2005).

### Limitations to Generalizability

While this research was able to successfully elicit the expected effects related to the ERP and spectral differences for the oddball and baseline tasks respectively, there are still several limitations of this study that must be further discussed. One such limitation is that we failed to quantify the participants’ perceived riding difficulty between the preferred and non-preferred conditions. This could have informed us about the degree of riding interference experienced by participants in the non-preferred condition and the relationship between subjective riding difficulty and overall ERP amplitude. Interestingly, Nenna et al. (2020) reported that even when participants do not experience subjective mental load in dual tasks when compared to a single task, it is still possible to obtain differences in neural responses. It could be possible that the expected increase in difficulty between the preferred and non-preferred conditions is being nullified by the electric skateboard in a way that participants can easily adapt to riding in their non-preferred stance since the board is self-propelling and very stable. We also did not measure torso muscle activity and acceleration on the body. Skateboarding has little overt movement but relies on balance using the torso and proximal muscles of the legs and arms. Further research could correlate changes in these muscle groups during different riding conditions with brain activity and sensory input to objectively quantify muscle involvement and motor task interference. More critically, by failing to record the acceleration between preferred and non-preferred conditions we could not record whether a hypothetical increase in task difficulty led to motor adjustments (e.g., slowing down) as a mechanism to enhance task performance (see Al-Yahya et al., 2011; De Sanctis et al., 2014). Additionally, while this skateboard paradigm offers the opportunity to greatly minimize covert movements (in comparison to walking or bicycle riding paradigms), it does not eliminate muscle movements methodically. Another limitation in the present study is the lack of an experimental condition where participants complete the oddball task while not moving. Our initial motivation to conduct this study was to test whether a change in riding difficulty leads to measurable changes in the ERP and oscillatory domains.

While the inclusion of a non-riding oddball condition would have improved the present results, we believe that the preferred vs non-preferred comparison is suitable for the present research question.

The auditory oddball task used did not result in significant behavioral, ERP, or spectral alpha power differences between conditions, contrary to our expectations. Given that the field of mobile EEG is developing, obtaining behavioral effects in line with ERP results is an important step in understanding brain processes in real-life scenarios. This limitation may be attributable to our sample size; however, given that we are able to measure the expected oddball and spectral alpha differences, with large effect sizes, we believe the lack of difference between our skateboarding conditions could be due to other limitations. The auditory oddball task we used may be too easy, with any behavioral or electrophysiological changes hitting a ceiling regardless of condition. A possible solution would be to utilize a more difficult auditory oddball task where the stimuli are more similar or introduce distractor stimuli throughout the task. Given that participants are on a constantly moving skateboard, with a constantly changing visual environment, a visual oddball task may be more appropriate and attentionally demanding.

## Conclusion

In the current study, we developed a skateboard EEG paradigm and replicated the classic oddball P3 effect while participants freely skated through a busy running track. While we found consistent target-standard tone differences in the P3 and MMN/N2b in frontal and parietal regions, we did not observe a reduction in P3 amplitude following increases in riding difficulty. We also present evidence that the classic peak in resting-state alpha completely diminishes while skateboarding and this overall desynchronization is the result of an increased load of incoming stimuli from the environment and task. The results in the current study support the notion that EEG paradigms are suitable for the study of human cognition under high ecological validity.

## Acknowledgments

This research was supported by start-up funds from the faculty of science and an NSERC Discovery grant (#RES0024267) awarded to Kyle Elliot Mathewson. The authors thank Pat Makhacheva for assistance in data collection, and Sarah Sheldon for analysis feedback.

## Author contribution

Robles: design, data acquisition, analysis/interpretation, drafting

Kuziek: design, data acquisition, analysis/interpretation, drafting

Wlasitz: design, data acquisition, drafting

Bartlett: study conception, design, data acquisition

Hurd: study conception, design, critical revisions

Mathewson: design, analysis/interpretation, drafting, critical revisions

## Data Accessibility Statement

This study’s supporting data and analysis material is open to access using the following link: https://github.com/APPLabUofA/Skateboard

## Abbreviations

ERP: Event-Related Potential,
EEG: Electroencephalography

